# An unbiased method to partition diverse neuronal responses into functional ensembles reveals interpretable population dynamics during innate social behavior

**DOI:** 10.1101/2024.05.08.593229

**Authors:** Alexander Lin, Cyril Akafia, Olga Dal Monte, Siqi Fan, Nicholas Fagan, Philip Putnam, Kay M. Tye, Steve Chang, Demba Ba, AZA Stephen Allsop

**Affiliations:** School of Engineering and Applied Sciences, Harvard University, Cambridge, Massachusetts, USA; Center for Brain Sciences, Harvard University, Cambridge, Massachusetts, USA; Kempner Institute for the Study of Artificial and Natural Intelligence, Harvard University, Cambridge, Massachusetts, USA; Department of Psychology, Yale University, New Haven, Connecticut, USA; Salk Institute for Biological Studies, La Jolla, California, USA; Howard Hughes Medical Institute, La Jolla, California, USA; Kavli Institute for the Brain and Mind, La Jolla, California, USA; Center for Collective Healing, Department of Psychiatry and Behavioral Sciences, Howard University, Washington DC, USA; Department of Psychiatry, Yale University, New Haven, Connecticut, USA

## Abstract

In neuroscience, understanding how single-neuron firing contributes to distributed neural ensembles is crucial. Traditional methods of analysis have been limited to descriptions of whole population activity, or, when analyzing individual neurons, criteria for response categorization varied significantly across experiments. Current methods lack scalability for large datasets, fail to capture temporal changes and rely on parametric assumptions. There’s a need for a robust, scalable, and non-parametric functional clustering approach to capture interpretable dynamics. To address this challenge, we developed a model-based, statistical framework for unsupervised clustering of multiple time series datasets that exhibit nonlinear dynamics into an *a-priori-*unknown number of parameterized ensembles called Functional Encoding Units (FEUs). FEU outperforms existing techniques in accuracy and benchmark scores. Here, we apply this FEU formalism to single-unit recordings collected during social behaviors in rodents and primates and demonstrate its hypothesis-generating and testing capacities. This novel pipeline serves as an analytic bridge, translating neural ensemble codes across model systems.

## Introduction

### Challenges in decoding neuronal representations during behavior

A common experimental approach in neuroscience involves recording neurophysiological activity from multiple neurons while a model organism engages in a specific task. The experimentalist must then understand how to use single neuron action potential data to decode the neural ensemble functions posited to represent stimuli and drive behavior ^1,2^. Conventional clustering methods, which depend on arbitrary categorization, pose challenges due to the variability in selecting the number of clusters and defining the criteria for each cluster. These approaches, therefore, are limited to coarse descriptions of population activity and do not capture changes in neural signals over time ^3,4^. Lastly, this process does not scale to large modern data sets containing hundreds to thousands of neuronal time series ^3,5^.

Automatically categorizing neurons into ensembles based on their response to an external stimulus can offer impartial insights into how neural representation functions computationally within groups of neurons across distributed networks. This process can also stimulate the formulation of biological hypotheses concerning how the brain encodes information. ^6^. As a result, various clustering methods have been designed to identify neurons by spike train patterns. They are grouped using computational methods that require the specification of predetermined parameters or numbers of groups ^7–9^. For example, nearest neighbors is a popular approach, but it requires training the model and for the experimentalist to define the size of the neighborhood, which can impact how data are clustered. Other modern approaches have fewer free parameters and are feature-based ^10^, lack interpretability because of their reliance on opaque artificial neural networks ^11, 12^, do not provide a direct link to the dynamics of a given ensemble ^13^, or are not based on state-space models of neural activity ^14^. State-space models can uncover the underlying neural dynamics with high efficiency and flexibility ^15^. Thus, there is a need for approaches that, starting with large amounts of time-varying neural activity, can not only cluster individual neurons into ensembles able to decode naturalistic behavior ^16,17^, but can also give interpretable insights into the nature of computations in these ensembles.

### A novel state-space-based unsupervised approach for clustering neurons into ensembles

Unsupervised approaches to clustering neural activity hold promise for solving the challenge of inferring the important features that define ensembles and using these characterizations to generate testable scientific hypotheses ^18^. To this end, state-space approaches are a compelling methodology to model neural population activity in a manner that can decode behaviors or cognitive states ^19–21^. Some of these methods collapse population data into a single state-space model ^22^, which treats all neural responses as part of a unified state process. This approach bypasses representations at the level of ensembles or single neurons. Our previous state-space method successfully modeled individual neuronal firing to generate novel insights about circuit function in social learning ^23^. However, this approach was limited by a lack of ability to define how individual neuronal firing contributed to ensemble representations during social learning.

To better understand ensemble activity, we recently derived a general, model-based, statistical framework for clustering multiple time series that exhibit nonlinear dynamics into an *a-priori*-unknown number of sub-groups or ensembles ^24^. In the context of neural data, we aim to use population neural spiking activity to automatically:

a. quantify the different types of neuronal responses, which we term Functional Encoding Units (FEUs)
b. characterize the dynamics that govern neuronal responses within each FEU by providing parameters with observable values.

We refer the reader to the Methods section of this manuscript for a brief mathematical explanation of our method for unbiased time-series clustering. Compared to previous feature-based clustering methods ^10^, this framework lets us perform direct statistical inference on the parameters of a physical model by which the neural time series are generated, such as a Hodgkin-Huxley model ^25^, an integrate-and-fire model ^25^, or point process state-space model ^19^. A Hodgkin-Huxley model defines neural activity in terms of the molecular components of the cellular membrane, such as calcium and sodium channel dynamics, while an integrate and fire model describes neural activity in terms of inputs that cross a certain activity threshold to generate an output. We utilize the point process state-space model because spike trains from multiple neurons exhibit multivariate point processes. This statistical framework offers flexibility and computational efficiency for neural decoding ^3,27^. Thus, FEU analysis provides an unbiased characterization of ensembles of neurons that allows novel insights at the level of population encoding that are not discernable from a single unit or whole population analysis. Each FEU ensemble can be formalized as a functional module ^28^, which can be widely distributed and variable, providing a substrate for representing the constantly fluctuating external and internal multi-modal stimuli that drive cognitive states and behavior.

### Using FEU representations to decode social information processing

Social behavior is dynamic and complex, and social representations are widely distributed across brain networks in various species ^29–32^. Social isolation can lead to pro-inflammatory gene expression, increased overall mortality ^33–35^, and has been linked to increased cardiovascular risk ^36^, functional decline ^37^, and increased likelihood of death ^38,39^. Lastly, some conditions, such as autism spectrum disorders and William’s syndrome, have particular social impairments ^40–42^. The relationship of social milieu to health is also seen in various other animals and is thus thought to be evolutionarily conserved and mediated by conserved neural mechanisms ^43,44^.

Given the evolutionary importance of social information processing to human health and the inherently distributed nature of social representations in the brain, we applied FEU analysis to neural data collected while rodents engaged in social learning ^23^ and primates engaged in live dyadic social gaze interaction ^45^. Social learning is a highly conserved social behavior with a distinct survival advantage. Previous work has demonstrated the conserved importance of the anterior cingulate cortex (ACC) and the basolateral amygdala (BLA) for representing social learning in rodents, primates, and humans ^46–49^. Eye gaze is a fundamental aspect of primate and human social behavior, and social gaze patterns effectively capture the underlying cognitive processes that are engaged during social interactions ^50^. Importantly, social gaze behaviors are also atypical in many psychiatric conditions, including autism ^51^, schizophrenia ^52^, and social phobia ^53^. Lastly, individual neurons in the ACC, BLA, dorsomedial prefrontal cortex (dmPFC), and orbitofrontal cortex (OFC) have been found to represent different social features during live social gaze ^45^.

FEU analysis applied to these datasets revealed novel ensemble dynamics occurring in these cortico-limbic regions in both species during complex forms of social behavior and demonstrates the hypothesis-generating and testing capacities of this analytical method.

## Results

### Simulation confirms accuracy of Functional Encoding Unit pipeline

We first demonstrate the accuracy of our FEU analysis pipeline by analyzing a simulated dataset of 50 noisy neurons belonging to 5 ground truth ensembles, each with different neural responses to a given stimulus: excited-sustained, excited-phasic, inhibited-sustained, inhibited-phasic, and nonresponsive (**Figure 1A**). In the pipeline, we used a state-space-GLM approach to model each simulated neuronal response as a linear state-space model ^23,54^. We performed the unsupervised clustering step by treating the population of neurons as a Dirichlet process mixture of state-space models and applying a Metropolis within-Gibbs clustering algorithm for inference (**Figure 1A**). Further technical details of the simulation, as well as a mathematical description of our mixture model, can be found in the Methods section.

**Figure 1.**
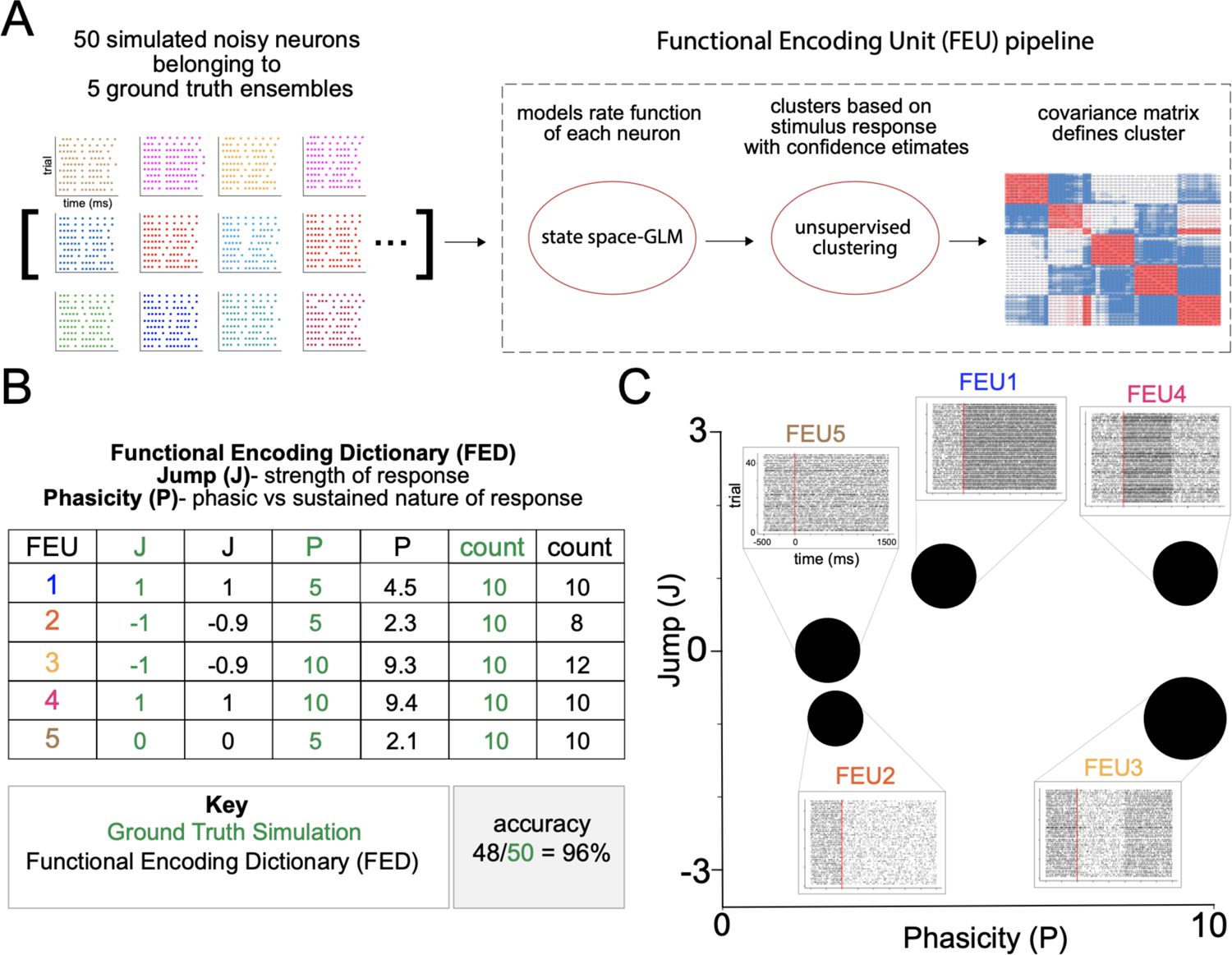
Functional Encoding Unit pipeline and validation. **A.** We combined a state-space approach for modeling neural firing with an unsupervised approach for clustering neurons based on their stimulus-response function. We simulated neural activity in a population of 50 noisy neurons belonging to five ground truth clusters of possible responses to a stimulus: excited-sustained, excited-phasic, inhibited-sustained, inhibited-phasic, and nonresponsive. To evaluate the accuracy of our approach for correctly identifying ensembles and making inferences about their parameters, we analyzed this simulated data using our Functional Encoding Unit (FEU) pipeline. **B.** The Functional Encoding Dictionary (FED) accurately clusters noisy neurons into the five ground truth ensembles: excited-sustained (FEU1), inhibited-sustained (FEU2), inhibited-phasic (FEU3), excited-phasic (FEU4), nonresponsive (FEU5) (48/50 = 96%). Each FEU has two parameters that define its function: the Jump (J) parameter and the Phasicity (P) parameter. J describes how much neuronal responses within an FEU are modulated by a given stimulus. A positive value indicates excitation, while a negative value indicates inhibition. A value close to zero (between −0.1 and 0.1) indicates no response. P describes how phasic or sustained the FEU response is to a given stimulus. A small value of P (< 5) indicates a sustained response, and larger values of P indicate a more phasic response. The FED provides parameter values for J and P that accurately model the ground truth relationships between FEUs. **C.** 2-D graphical representation of FED. FEUs have centroids corresponding to their value of J and P parameters; the size of the FEU represents the relative number of neurons in each FEU. Callout to rasters for each FEU made from the individual rasters of all neurons in that FEU.

The FEU pipeline demonstrated high accuracy (48/50 neurons, 96%) in clustering the 5 ground truth ensembles into 5 FEU and correctly characterized each FEU with two parameters defining the relative ensemble response: Jump (J) and Phasicity (P) (**Figure 1B, C**). J describes how much the response of neurons within an ensemble is modulated by a given stimulus. A positive value indicates excitation, a negative value indicates inhibition and a value close to zero (between −0.1 and 0.1) indicates no response. The second parameter, P, describes how phasic or sustained an ensemble response is to a given stimulus ^23^. A small value of P (less than 5) indicates a sustained change in response to the stimulus, whereas larger values of P indicate that the change is phasic or unsustained (**Figure 1B**). The number of FEUs, the value of their J and P parameters, and the number of neurons within each FEU comprise our Functional Encoding Dictionary (FED), in which each FEU can be plotted into a 2-dimensional space to visualize the population ensemble representation (**Figure 1C**). In the graphical space, each FEU has a centroid located at its parameter values, and the size of the FEU is relative to the number of neurons within it (**Figure 1C**). The five ground truth ensembles were separable in FEU space, and the FED representation recovered the total ensemble representation within the simulated population of neurons (**Figure 1C**). Lastly, peri-event rasters were constructed from the action potentials of all neurons within each FEU, demonstrating the captured ensemble dynamics (**Figure 1C**).

### FEU provides interpretable latent space and outperforms other clustering methods

We compared the latent space and cluster analysis learned by FEU to those of other methods in the literature. Latent spaces are often uninterpretable or unintuitive due to the black-box nature of the machine-learning algorithms that produce them. When dealing with complex data, dimensions in the latent space may not correspond to meaningful properties of the data, posing challenges for interpretation. To generate quantitative metrics for clustering performance, we conducted multiple simulations where the ground-truth clusters were known. K-Means is an established methodology for clustering data, though it has the limitation of requiring the user to specify the number of clusters K ^55–57^. A popular approach in the literature for clustering high-dimensional data is to (1) use a dimensionality reduction technique (e.g., PCA, T-SNE, CEBRA) to map data to a latent space with fewer dimensions and (2) use K-Means to cluster the data in the latent space ^58–64^. Following this structure, we compared FEU against several techniques, including K-Means on the raw data, PCA + K-Means, T-SNE + K-Means, and CEBRA + K-Means. We refer the reader to the Methods section for further methodological details for this evaluation. K-Means clustering analysis showed incorrect clustering of the data (**Figure 2A**). PCA + K-Means and T-SNE + K-Means show slightly better clustering but are still mostly incorrect (**Figure 2B, 2C**). Among the benchmark methods, CEBRA + K-Means clustering analysis showed the most accurate clustering but was outperformed by the FEU pipeline (**Figure 2D, 2E**). The accuracy of the FEU pipeline can be highlighted by the fact that most neurons overlapped in the FEU space and required a callout to visualize all neurons (**Figure 2E, Supplementary** Figure 1).

**Figure 2.**
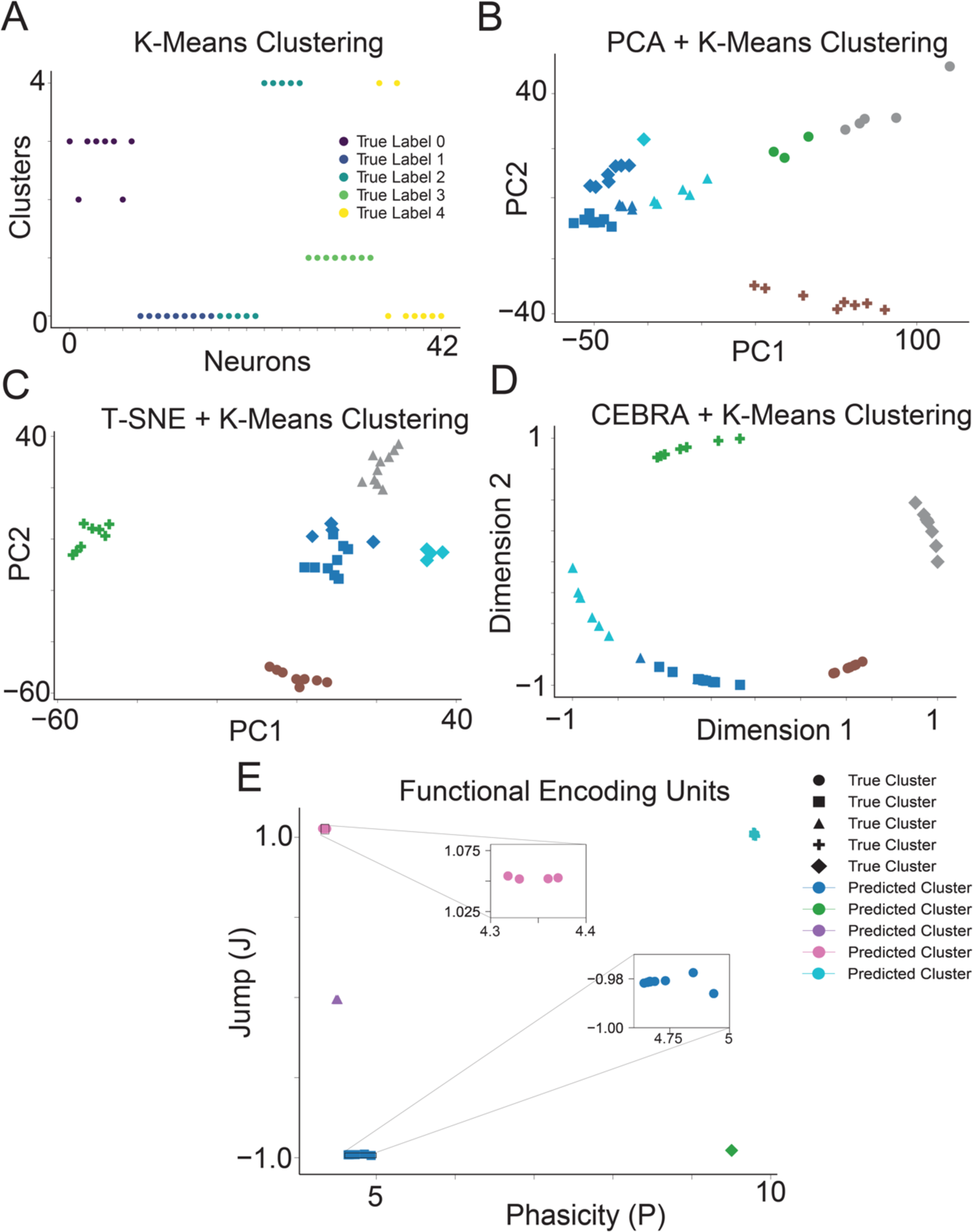
FEU analysis clusters simulated data more accurately than other methods. **A.** We applied K-Means to cluster simulated neuron spike data. Data points correspond to individual neurons; the y-axis ‘Clusters’ denotes each neuron’s cluster assignments determined by K-Means, with colors indicating the neurons’ true clusters. Perfect clustering is achieved when neurons within the same predicted cluster share identical colors, reflecting accurate alignment with their true cluster. **B.** We performed dimensionality reduction on the simulated neuron spike data with PCA and clustered the resulting principal components with K-Means. Data points correspond to individual neurons. The data point shape denotes the ground truth cluster assignment, while colors denote the predicted clusters. Perfect clustering is achieved when data points in clusters have the same color and shape. **C.** We conducted dimensionality reduction on the simulated neuronal spike data using T-SNE, followed by clustering the derived principal components with K-Means. Each data point represents an individual neuron, with the shape of the data point indicating the neuron’s actual cluster assignment and colors representing the predicted clusters. Perfect clustering is achieved when data points in clusters have the same color and shape. **D.** We performed dimensionality reduction on the simulated neuronal spike data utilizing CEBRA and subsequently clustered the resultant components with K-Means. Each data point signifies an individual neuron; data point shape denotes the ground truth cluster assignment, while colors denote the predicted clusters. Perfect clustering is achieved when data points in clusters have the same color and shape. **E.** FED representation of the simulated neuron spike data. We use FEU analysis to cluster the simulated neuron spike data. Perfect clustering is achieved when data points in each cluster have the same color and shape. Callout to show overlapping data points in a cluster.

To quantify the performance of our algorithm in relation to other methods, we used normalized mutual information score (NMI), adjusted rand index (ARI), and accuracy. NMI quantifies the degree of overlap in information shared among clusters. An NMI value of 1.0 signifies optimal clustering, while 0 suggests ineffective clustering. ARI measures the agreement between data partitions ^62,65^ and is normalized to account for randomness. An ARI value of 1 indicates that the clustering is identical, while a value close to 0 suggests random clustering and negative values indicate independent or dissimilar clustering ^66^. Accuracy in this context is the percentage of correctly predicted labels after the alignment of cluster labels between the true and predicted labels using the Hungarian algorithm.

There were significant differences in performance between the FEU pipeline and the other benchmark methods on our multiple simulations (**Figure 3).** FEU outperforms K-Means clustering, PCA + K-Means, T-SNE + K-Means, and CEBRA + K-Means in all evaluation metrics with less variability in scores across multiple simulations (**Figure 3**), despite the fact that the optimal number of clusters K was always supplied to K-Means whereas FEU had to learn the number of clusters directly from the data. FEU pipeline achieved an NMI score of 0.977 ± 0.024 (**Figure 3A**), demonstrating a high degree of overlap between the true cluster labels and the predicted labels, an ARI score of 0.973 ± 0.03 (**Figure 3B**), showing that the clustering closely matches the true cluster labels, and lastly, an accuracy score of 0.979 ± 0.019 (**Figure 3C**), which means that a higher proportion of points where correctly clustered. These results show that the FEU pipeline is robust, and its clustering is more aligned with the true underlying structure of the data.

**Figure 3.**
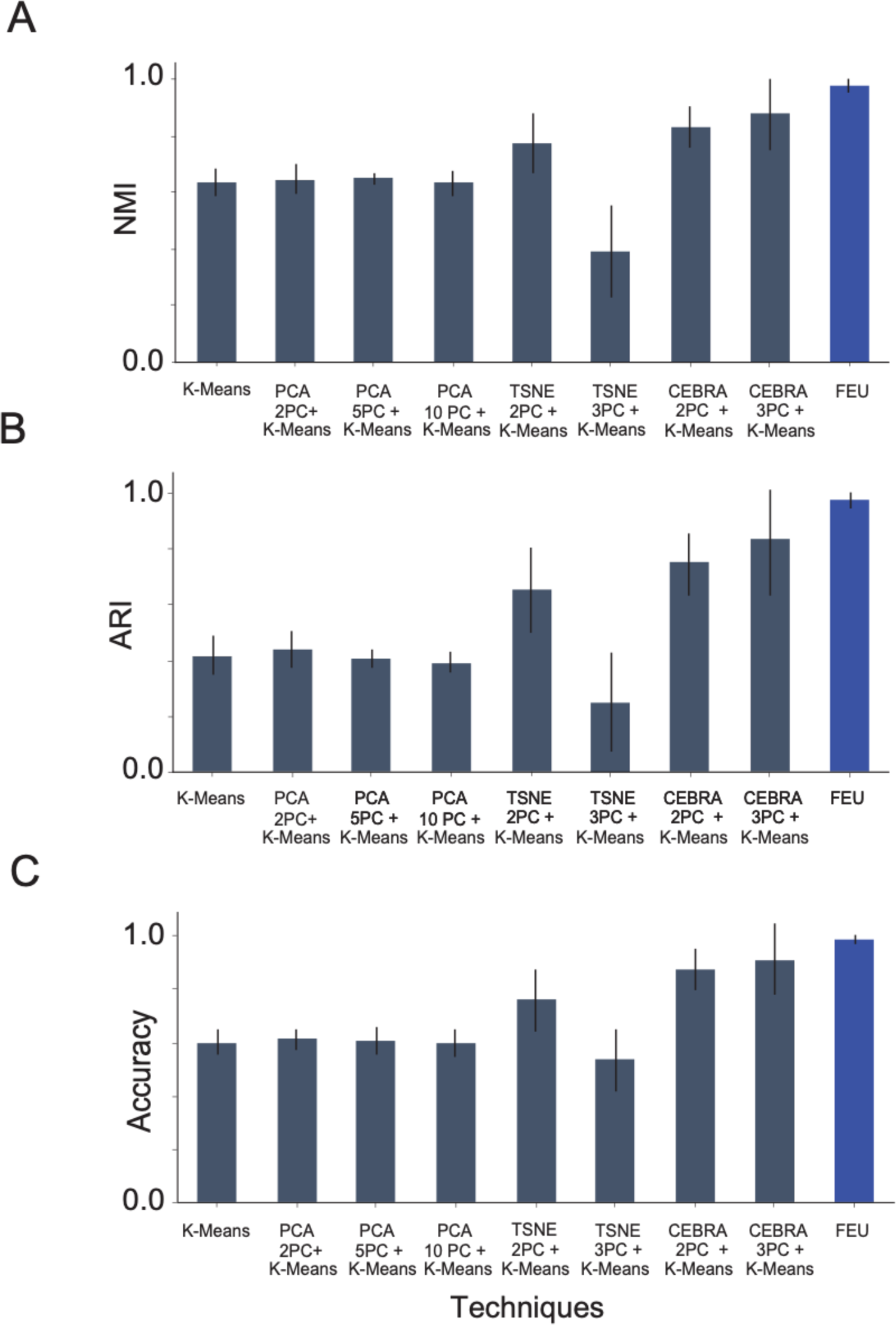
FEU analysis outperforms other methods over multiple simulated datasets. **A.** We compare the NMI score of FEU analysis to alternative methods when tested over five experiments of simulated data. The alternative methods are K-Means, PCA + K-Means (using 2, 5, and 10 principal components), and T-SNE + K-Means (using 2 and 3 principal components). The presented score is the mean NMI score across five datasets with error bars denoting the standard deviation. FEU pipeline achieved an NMI score of 0.977 ± 0.024 **B.** The ARI score of FEU analysis is compared to alternative methods when tested over 5 experiments of simulated data. The alternative methods are K-Means, PCA + K-Means (using 2, 5, and 10 principal components), and T-SNE + K-Means (using 2 and 3 principal components). The presented score is the mean ARI score across five datasets with error bars denoting the standard deviation. FEU pipeline achieved an ARI score of 0.973 ± 0.03 **C.** The accuracy of FEU analysis in comparison with alternative techniques across five simulated data experiments is demonstrated. These techniques include K-Means, PCA + K-Means (using 2, 5, and 10 principal components), and T-SNE + K-Means (using 2 and 3 principal components). The reported score represents the average accuracy obtained from these five datasets, highlighted with error bars to indicate the standard deviation. FEU pipeline achieved an accuracy score of 0.979 ± 0.019

Finally, we observe that the latent space learned by FEU has axes that can be mapped to model parameters with a physical interpretation in terms of the intensity function. In contrast, the latent spaces of other methods, such as PCA, T-SNE, and CEBRA, are not directly interpretable because they generally find directions of maximal data variance, which may or may not be aligned with physical quantities of interest.

### FEU analyses of neural activity recorded while rodents engaged in social learning

Given the model’s performance on simulated data, we applied FEU analysis to in vivo action potential data recorded from neurons in the anterior cingulate cortex (n=195) and the amygdala (n=72) of rodents engaged in social learning ^23^ (**Figure 4A**). During social conditioning, a cue predicts shocks delivered to a demonstrator mouse (d). We recorded from neurons in the ACC or BLA of the observer mouse (o) who learned the predictive value of the cue (**Figure 4A**). FEU analysis clustered ACC and BLA activity into 9 FEU ensembles (**Figure 4B**). FEUs represented a range of responses that include excited, inhibited, sustained, and phasic responses (**Figure 4B, 4C)**.

**Figure 4.**
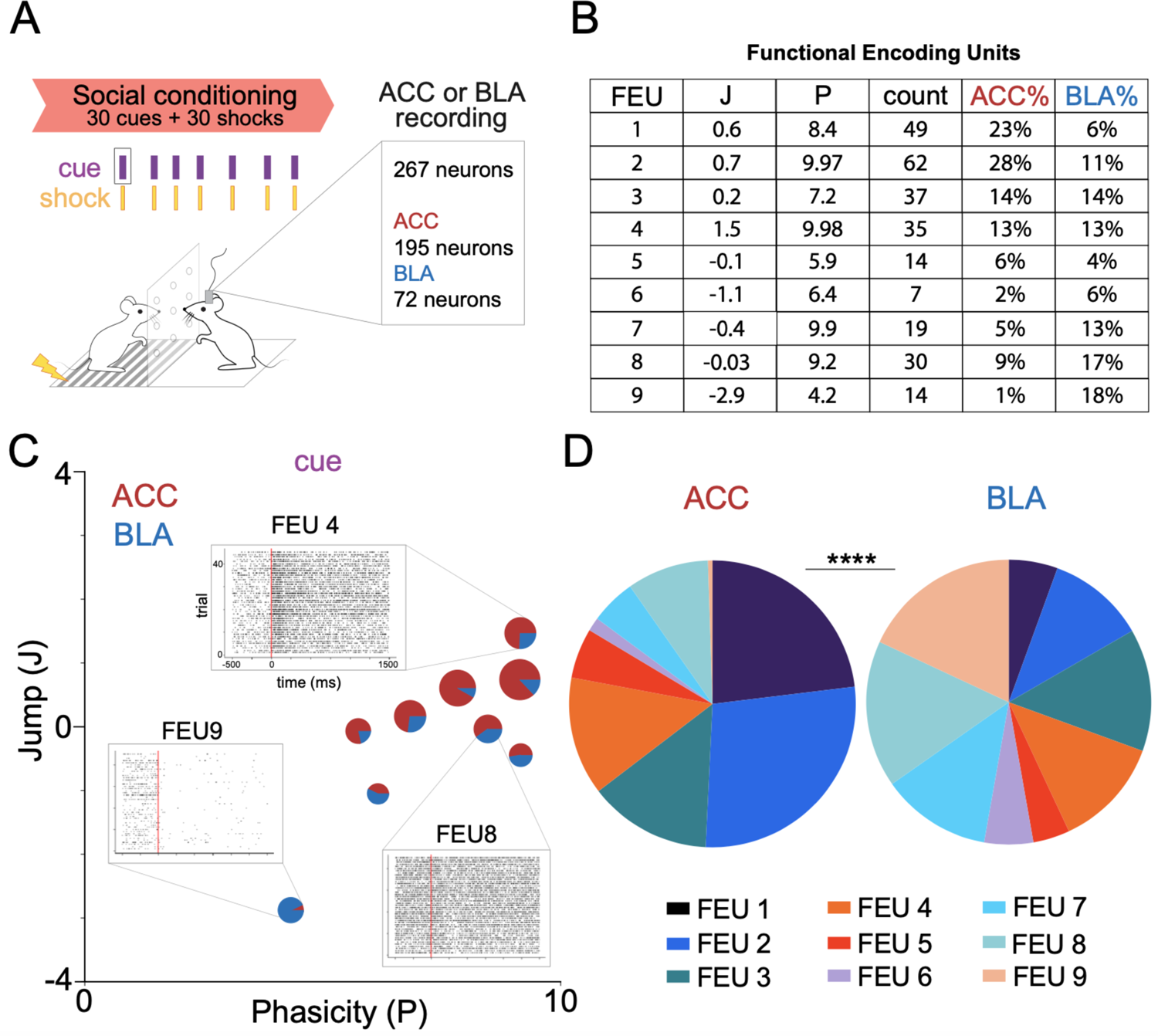
ACC and BLA have shared FEU representations during social conditioning. **A.** We applied FEU analysis to neurons recorded from the anterior cingulate cortex (ACC) and basolateral amygdala (BLA) of mice engaged in a social conditioning paradigm in which observer mice (o) learn that a cue predicts delivery of a shock to a demonstrator (d) mouse (Allsop et al. 2018). **B.** FED reveals 9 FEU ensembles with neurons from the ACC and BLA in each ensemble. Jump (J) and Phasicity (P) values define FEU ensemble responses to the cue during social conditioning. The count represents the number of neurons in each FEU, while the ACC% and BLA% are the percentage of the recorded ACC or BLA neurons, respectively, that are in each FEU. **2**-D graphical FED representation of ACC and BLA FEU cue activity during rodent social conditioning. Each FEU is composed of varying proportions of both ACC and BLA neurons. Call out to representative FEU rasters illustrative of varying response dynamics captured by FEU ensembles: excited and phasic (FEU 4), non-responsive (FEU 8), inhibited and sustained (FEU 9). ACC and BLA have different proportions of their neurons in cue FEU ensembles during social conditioning. Pie charts represent all neurons recorded from each region and the proportion of neurons recorded from each region that belong to each FEU (Chi-square test; χ^2^ = 55.97, df= 8, ****p<0.0001).

Interestingly, each FEU contained neurons from both the ACC and BLA (**Figure 4B, 4C**). Given previous findings demonstrating the structural and functional connection between the ACC and BLA during social learning ^23,67–70^, the finding that the ACC and BLA co-cluster into functional ensembles has external validity with existing literature. Notably, however, the proportion of neurons from the ACC and BLA that cluster into specific FEUs was significantly different between the two regions (**Figure 4D**), suggesting some discernible separation in their FEU code (Chi-square test; χ^2^ = 55.97, df= 8, ****p<0.0001).

.We also applied CEBRA + K-Means and K-Means clustering to the ACC and BLA neural data (**Supplementary** Figure 2). Though CEBRA + K-Means identifies some clusters with meaningful structures, it does not capture the diversity of raster structures identified by FEU; for example, CEBRA + K-MEANS failed to find certain notable patterns (e.g., neurons with inhibited firing dynamics) **(Supplementary** Figure 3).

We previously found that neurons within the ACC demonstrate functional plasticity and change their firing rate to a cue during social learning ^23^. We tested whether FEU analysis could capture neural dynamics that occur during social learning. We compared FEU ensemble representations of the cue during the habituation period of social learning when the cue had no predictive value to the social conditioning period when the cue predicted delivery of shock to the demonstrator (**Figure 5A, Table 1**). To ensure that FEU ensemble clusters were determined by representations of the cue, we simulated a reference event (simulated cue) in which there was no external stimulus delivered and clustered FEU activity around this “simulated cue” as a control analysis (**Figure 5A-D**). FEU cue representations within the observer’s ACC shifted during social learning, with FEUs having more sustained firing (P<5) during habituation and more phasic firing (P>5) during conditioning (**Figure 5B**). Notably, the average Jump of all FEUs was not different between conditions (Welch’s ANOVA; W (DFn, DFd) = 0.01439 (3.000, 15.86) p= 0.9975). (**Figure 5C**). However, the average Phasicity during cue representation significantly increased during social conditioning when compared to the control condition, suggesting that increased Phasicity of ensembles may encode learning (Welch’s ANOVA; W (DFn, DFd) = 0.01439 (3.000, 15.86) p= 0.9975) (**Figure 5D**).

**Figure 5.**
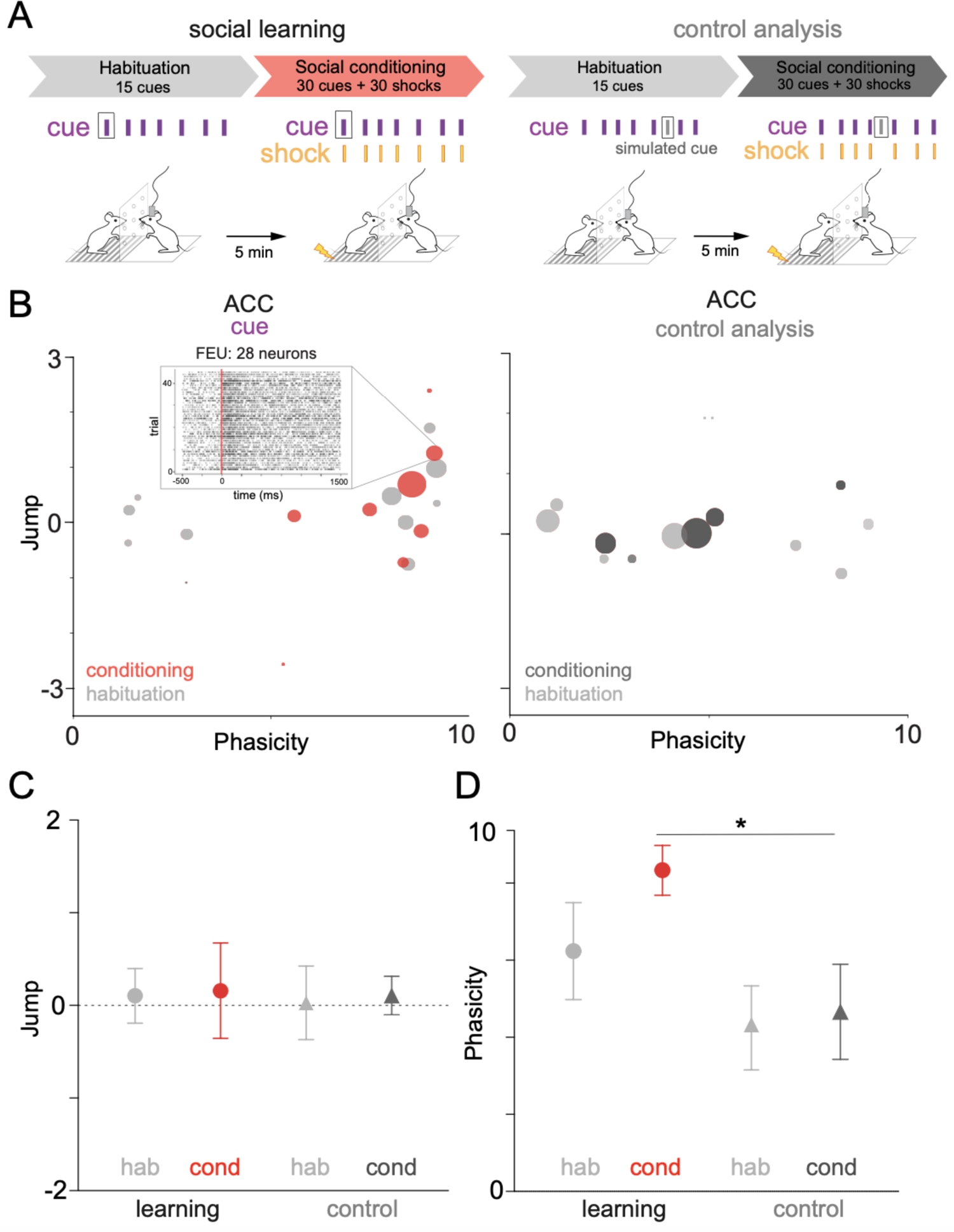
FEU representations capture changes in neural activity that occur with social learning. **A.** We applied FEU analysis to the same social conditioning paradigm while including a habituation period in which cues were not predictive of shock delivery. We could then compare neural responses during habituation to those during social conditioning when cues predict shock delivery to the demonstrator. To test whether FED representations capture information about the cue, we included a control condition in which the pipeline received a false cue input (control analysis). **B.** 2-D graphical FED representation of ACC cue FEU activity during habituation and conditioning periods of rodent social learning (left). Control analysis showing a FED representation of the false cue during habituation and conditioning (right). Call out to raster of selected FEU with 28 neurons and an excited-phasic cue response during social conditioning. See Table 1 for FED values. **C.** Average cue Jump of all ACC FEUs during habituation and conditioning of social learning. No significant difference was found between the average ACC cue Jump during habituation and conditioning (Welch’s ANOVA; W (DFn, DFd) = 0.01439 (3.000, 15.86) p= 0.9975). **D.** Average cue Phasicity parameter of all ACC FEUs during habituation and conditioning of social learning. There is a significant difference between FEU cue Phasicity during conditioning when compared to the control condition (Welch’s ANOVA; W (DFn, DFd) = 4.092 (3.000, 13.82), *p< 0.0283, Dunnett’s post-test, learning-cond vs control cond, *p = 0.0385).

A benefit of this statistical approach is that it can infer the properties of a biophysical model of neural activity. We previously described a dual-viral, optogenetic approach allowing for experimental phototagging of three ACC-BLA networks with known anatomic or biophysical features ^23^. Neurons in the photoidentified network directly project from the ACC to the BLA (**Figure 6A**). Excited network neurons have reciprocal or collateral inputs from photoidentified neurons (**Figure 6A**). Inhibited network neurons have reciprocal or collateral inhibitory inputs from the photoidentified network (**Figure 6A**). We applied the known network identities of neurons to the previously clustered FEUs during social learning (**Figure 6B, Table 2**). We tested the ability of FEU analysis to differentiate between anatomically and functionally distinct ACC-BLA networks during social learning (**Figure 6C-E**). Photoidentified, excited, and inhibited network neurons showed distributed representations across the FEU ensembles during habituation and conditioning (**Figure 6C**). However, ACC photoidentified and excited networks had higher average cue Jump values than the inhibited network (Welch’s ANOVA; W (DFn, DFd) = 5.911 (5.000, 42.58), ***p< 0.0003, Dunnett’s post-test; excited-hab vs inhibited-hab, **p = 0.0030; excited-cond vs inhibited-cond, *p = 0.0286) (**Figure 6D**). Interestingly, the inhibited network also had lower average cue Phasicity values during habituation than the photoidentified network and exhibited a significant increase in Phasicity during social learning (Welch’s ANOVA; W (DFn, DFd) = 4.499 (5.000, 43.45), **p=0.0022, Dunnett’s post-test; inhibited-hab vs inhibited-cond, **p = 0.0194). (**Figure 6E**). Thus, the average parameters that define ACC cue FEU representations were significantly different across biophysically distinguishable ACC-BLA neural networks (i.e., photoidentified vs inhibited network) during social learning. Additionally, FEU representation suggests that changes in ensemble representation during learning may be driven by changes in the inhibitory network.

**Figure 6.**
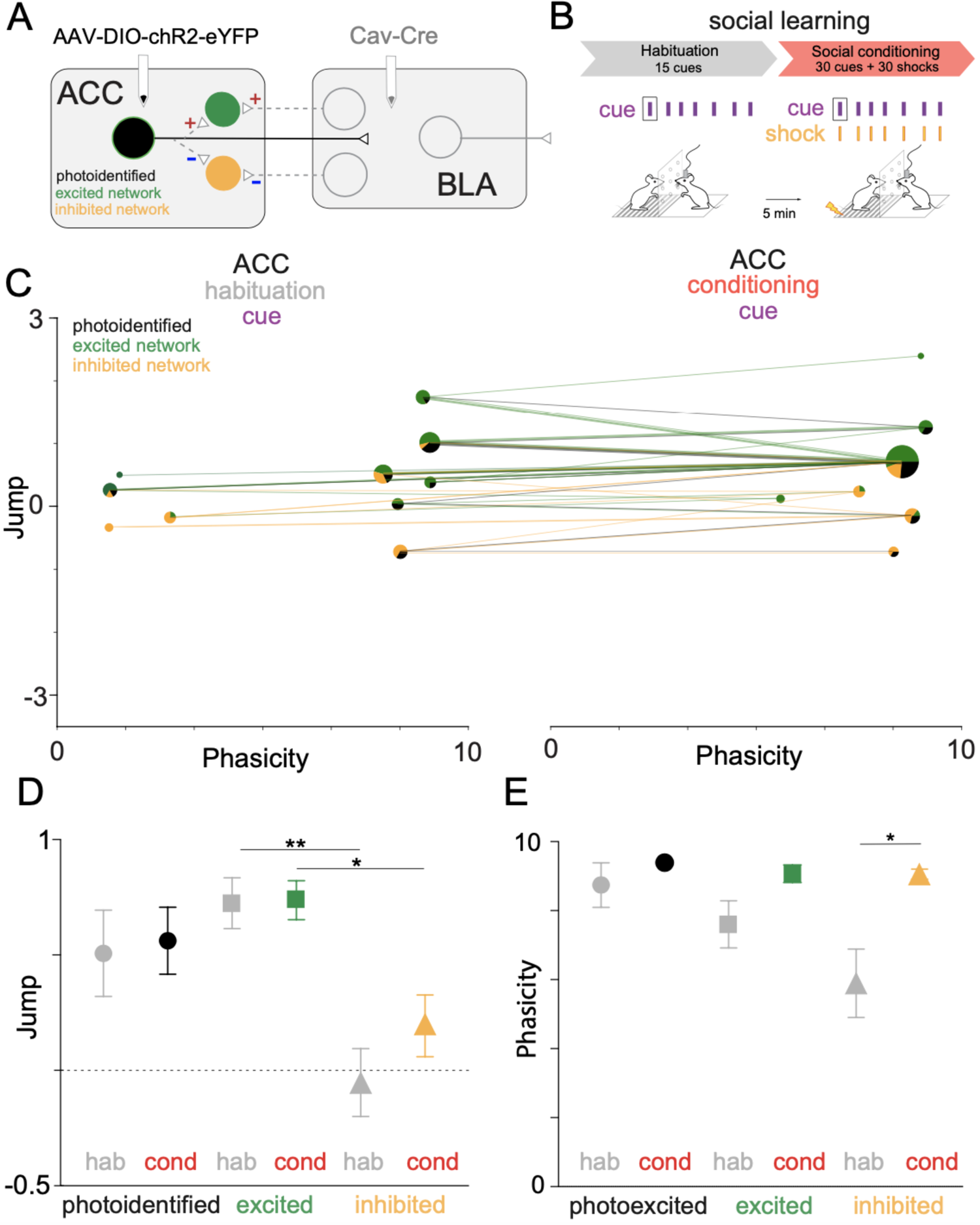
ACC-BLA networks have different cue FEU representations during social learning. **A.** An optogenetic approach allowed the targeting of neuronal circuits via direct ACC-BLA projection stimulation. We injected retrograde virus CAV2-Cre into the BLA and AAV-DIO-ChR2-eYFP into the ACC to express ChR2 in ACC neurons that monosynaptically project to the BLA. We identified network connectivity of ACC neurons based on in vivo phototagging, revealing three light-responsive categories: photoidentified, ACC→BLA excited, or ACC→BLA inhibited network. **B.** We informed our FEU analysis during habituation and conditioning of social learning with the network identity of ACC neurons. **C.** 2-D graphical FED representation of ACC-BLA network cue FEU activity during habituation (left) and conditioning (right) periods of rodent social learning. FEU ensembles contain varying proportions of photoidentified (direct ACC→BLA projectors), excited network, and inhibited network neurons. Lines show the trajectory of individual neurons between FEUs as ensemble activity changes from habituation to conditioning during learning. **D.** Average cue Jump parameter for photo-identified, excited, and inhibited networks during habituation and conditioning of social learning. Neurons in excited networks have higher average cue Jump parameters than inhibited network neurons during habituation and conditioning (Welch’s ANOVA; W (DFn, DFd) = 5.911 (5.000, 42.58), ***p< 0.0003, Dunnett’s post-test; excited-hab vs inhibited-hab, **p = 0.0030; excited-cond vs inhibited-cond, *p = 0.0286). **E.** Average cue Phasicity for photoidentified, excited, and inhibited networks during habituation and conditioning of social learning. The inhibited network shows a significantly higher average cue Phasicity during the conditioning period of social learning than habituation. (Welch’s ANOVA; W (DFn, DFd) = 4.499 (5.000, 43.45), **p=0.0022, Dunnett’s post-test; inhibited-hab vs inhibited-cond, **p = 0.0194).

### FEU analyses of primate neural activity demonstrate the versatility of the FEU formalism

Given its state-space approaches, FEU utilizes a unifying framework for analyzing neural ensemble representations that may be translatable across behavioral contexts as well as model organisms. We, therefore, probed the capacity of the FEU approach to cluster neurons in richer datasets such as those generated by recordings in non-human primates involved in naturalistic social behaviors. We applied FEU analysis to data collected during naturalistic social gaze interaction between pairs of rhesus macaques ^45,71^. Monkeys were positioned face-to-face and were able to spontaneously interact using gaze while the eye positions of both monkeys were tracked continuously at high resolution (**Figure 7A**). During these social gaze interactions, in vivo, action potential data was recorded from the anterior cingulate cortex (ACC), basolateral amygdala (BLA), dorsomedial prefrontal cortex (dmPFC), or orbitofrontal cortex (OFC) of monkeys (**Figure 7B**). Here, we focused on comparing FEUs when the recorded monkey looked at the partner’s face versus a non-social object (**Table 3**). ACC face ensembles were largely distinct from object ensembles during live social gaze (**Figure 7C**), and the average ACC face J had a decreased variance compared to the average J during object representation (Welch’s t-test; t=1.038, df=5.631, p=0.3418; F, (DFn, Dfd) = 18.76, (5, 4), *p=0.0140) (**Figure 7C inset**).

**Figure 7.**
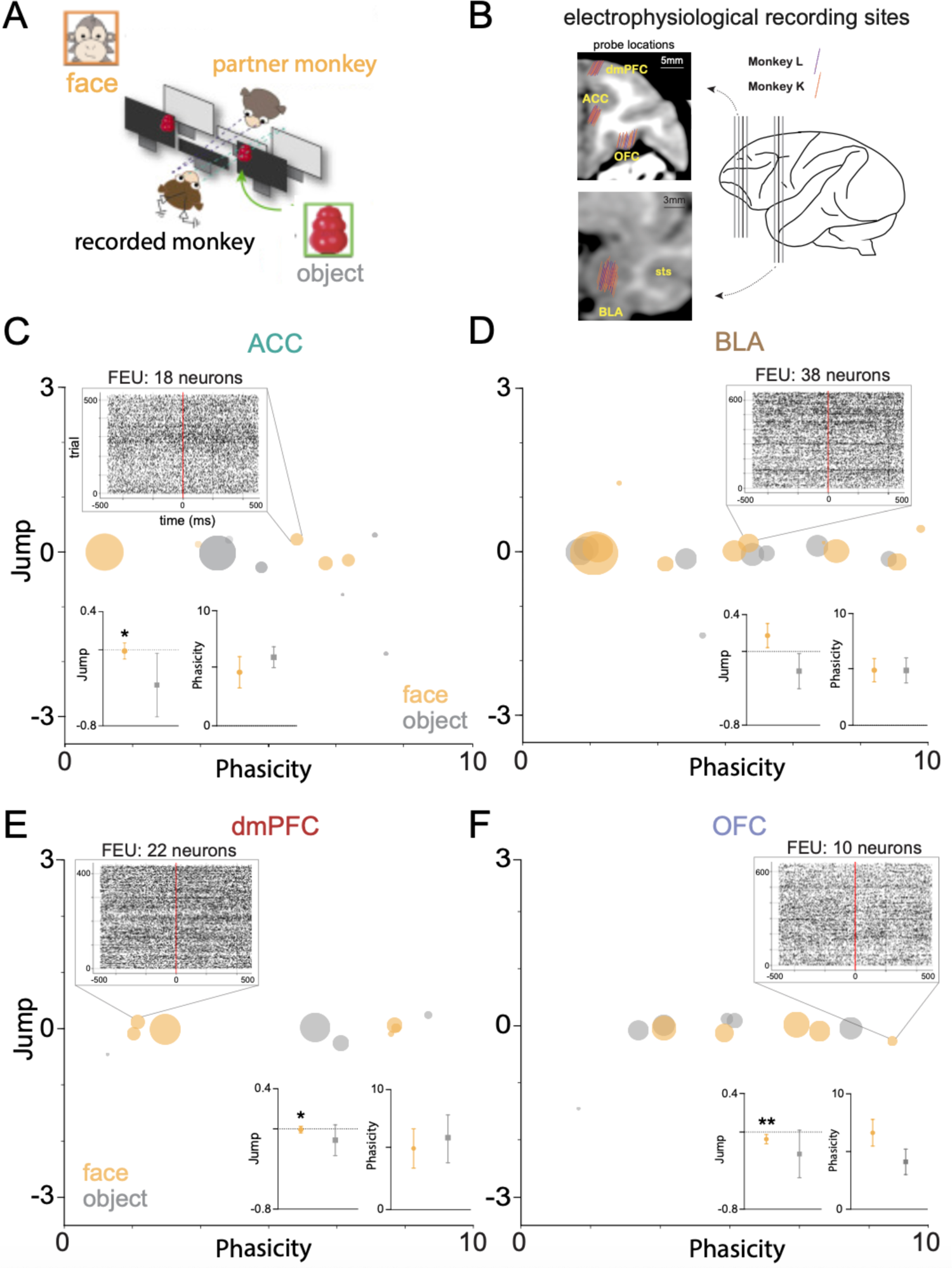
Face and Object FEDs across four non-human primate cortico-limbic regions. **A.** We applied FEU analysis to data from Dal Monte et al. 2022, in which rhesus macaque monkeys interact during a live social gaze interaction task. A freely visible object is also placed within the visual range of the monkey. Eye tracking data is collected to determine time stamps for when the recorded monkey looks into its partner’s face versus the nearby object. **B.** Probe locations for in vivo electrophysiological signal recordings from neurons in the anterior cingulate cortex (ACC), Basolateral amygdala (BLA), dorsomedial prefrontal cortex (dmPFC), or orbitofrontal cortex (OFC) of rhesus macaques L and K during the live social gaze paradigm. **C.** 2-D graphical FED representation of ACC face and object FEU activity during live social gaze. Call out shows a raster of representative FEU composed of 18 neurons and an excited-phasic response to a partner monkey’s face. Insets show average Jump and Phasicity during face vs object representation. ACC shows decreased variance in average face Jump parameter during social gaze compared to object gaze (Welch’s t-test; t=1.038, df=5.631, p=0.3418; F, (DFn, Dfd) = 18.76, (5, 4), *p=0.0140). **D.** FED representation of BLA face and object FEU activity during live social gaze. Call out shows a raster of representative FEU composed of 38 neurons and an excited-sustained response to a partner monkey’s face. Inset shows average Jump and Phasicity during face vs object representation. There was no significant difference in the Jump parameter (Welch’s t-test; t=0.6752, df=3.2379, p=0.1196; F, (DFn, Dfd) = 1.708, (7, 9), ns). **E.** FED representation of dmPFC face and object FEU activity during live social gaze. Call out shows a raster of representative FEU composed of 22 neurons and an excited-sustained response to a partner monkey’s face. Inset as above. dmPFC shows decreased variance in average face Jump across FEUs during face representation (Welch’s t-test; t=0.6752, df=12.89, p=0.5441; F, (DFn, Dfd) = 14.45, (3, 5), *p=0.0135) **F.** FED representation of OFC face and object FEU activity during live social gaze. Call out shows a raster of representative FEU composed of 10 neurons and an inhibited-phasic response to a partner monkey’s face. Inset as above. OFC shows decreased variance in average Jump across FEUs during face representation (Welch’s t-test; t=0.6<u>105</u>, df=5.375, p=0.5664; F, (DFn, Dfd) = 31.77, (5, 4), **p=0.0052)

By contrast, BLA face and object ensembles were closely associated in FEU space and showed no average parameter differences. However, there was a trend for an average higher J during face representation (Welch’s t-test; t=0.6752, df=3.2379, p=0.1196; F, (DFn, Dfd) = 1.708, (7, 9), ns) (**Figure 7D inset**). Similar to the ACC, dmPFC face ensembles were distinct from object ensembles (**Figure 7 E**), and the average dmPFC J had a decreased variance during face representation compared to object representation (Welch’s t-test; t=0.6752, df=12.89, p=0.5441; F, (DFn, Dfd) = 14.45, (3, 5), *p=0.0135) (**Figure 7E inset**). Lastly, OFC face and object ensembles were closely associated in FEU space (**Figure 7F**), and like the ACC and dmPFC, the average OFC Jump had decreased variance during face representation (Welch’s t-test; t=0.6<u>105</u>, df=5.375, p=0.5664; F, (DFn, Dfd) = 31.77, (5, 4), **p=0.0052) (**Figure 7F inset**). Thus, FEU characterization of the 4 regions during live social gaze interaction captured varying representations between the three cortical regions (ACC, dmPFC, OFC) and the subcortical region (BLA).

## Discussion

As the brain is likely to represent stimulus information through coordinated neural activity, it’s crucial to understand how distinct aspects of stimuli are encoded by distinct subsets of neurons sharing common encoding properties. We developed a neural data analysis pipeline that learns the number of distinct response profiles within a population of neurons and partitions the population into distinct apriori unknown ensembles (FEUs), each characterized by a stereotypical response profile defined with parameters. Together, the number of FEUs, the response parameters of each ensemble, along the biological properties of the neurons within the ensemble make up a Functional Encoding Dictionary (FED) (**Table 1,2,3**), a novel rich representation of ensemble encoding for a given brain region.

First, our pipeline outperformed current state-of-the-art techniques on simulated ground truth data and had high accuracy in recovering the correct number of ensembles and determining the correct relative responses of the ground truth ensembles (excited-phasic, excited-sustained, inhibited phasic, inhibited sustained, nonresponsive). In contrast to current state-of-the-art approaches, FEU analysis does not require *a priori* selection of the correct number of ensembles as it infers this correctly from the data structure itself. Additionally, it captures a broader range of ensemble responses than other techniques. Importantly, in contrast to other contemporary methods, FEU’s latent space is also mapped to parameters with biophysical interpretation that can be defined by an intensity function.

During social conditioning, FEU analysis indicates that neurons in the ACC and BLA are functionally connected during a task such as social learning, and these neurons from both regions are co-distributed across the same ensembles rather than cluster into independent ensembles (**Figure 4**). This suggests that neurons from anatomically connected regions form functional ensembles to achieve distributed neural representation during social behavior.

FEUs also captured changes in ensemble representations that occur during social learning and, importantly, found an increase in the average cue’s Phasicity during social learning (**Figure 5**). Previous experiments have shown that optogenetically induced phasic firing of dopamine drives reward learning, while sustained firing does not ^72^. Other experimental work demonstrates that more phasic populations have a stronger linear response to increases in synaptic input and are less dependent on background activity levels ^73^. Thus, there are biologically plausible mechanisms to relate observed changes in FEU Phasicity to learning. Our approach allows for future experimental work that can directly test how changes in measurable biophysical properties of synaptic physiology lead to changes in the Phasicity parameter.

We also found that labeling the network identity of neurons revealed significant average differences in FEU parameters between the networks (**Figure 6**). Particularly, we found that the inhibited network showed the greatest change in Phasicity from habituation to conditioning and is related to the ACC-BLA direct projectors through GABAergic signaling. Our analysis highlights a novel role of the ACC-BLA inhibitory network signaling during social learning. The importance of inhibitory network signaling in social learning is predicted by computational models of cortical neural population coding ^74^. It is also consistent with data from genetic experiments showing that optogenetic control of GABA somatostatin neurons in the ACC modulates social learning ^75^. Thus, FEU parameters reflect tangible neurobiological properties governing the evolution of ensemble representation, and FEDs may shed light on how network identity sets the constraints for the response profiles of ensembles in the brain. Future research focusing on recording from thousands of genetically identified neurons across multiple brain regions and employing FEDs as real-time outputs will elucidate the direct relationship between FEU parameters and network computations. Forthcoming experiments will directly investigate how processes such as Long-Term Potentiation (LTP) influence parameters like Jump, which is hypothesized to depend on both the number and strength of inputs to an ensemble. Optogenetic or electrical stimulation can also be utilized to directly examine the impact of different oscillations within ensembles on Phasicity.

Our FEU cluster assignment models the dynamic responses of ensembles of neurons during a specific stimulus. Network-level interactions between neurons are implicit in the fact that neurons that belong to the same cluster exhibit similar dynamics and, hence, covary. A limitation of this current approach is that it does not explicitly model the dependence of a given neuron’s response on the responses of other neurons in the network ^76^. It remains unknown what network population model is consistent with a mixture of state-space models as introduced here.

There is a large translational gap in neuroscience and psychiatry, partially due to a lack of analytical tools that span model systems and behavioral paradigms. We demonstrated here that FEU analysis can be applied to non-human primate data and is able to reveal different representational strategies across cortico-limbic networks involved in live social gaze **(Figure 7).** Notably, face FEU ensembles in the ACC, dmPFC, and OFC had a lower variance in the Jump parameter when compared to the object FEUs. This suggests more uniform Jump parameters in cortical regions for social stimuli, or stimuli that are more salient compared to the object. This interpretation is supported by previous descriptions of decreased response variability to a preferred stimulus across intracellularly recorded cortical neurons ^77^. Additionally, decreased variance of FEUs in cortical regions may decrease noise in the population signal, leading to better stimulus representation ^78^. Future work will further characterize the mechanisms that result in the observed changes in the variance of FEU parameters and how variance in individual neurons contributes to these changes ^79^.

The data-driven, unsupervised nature of our approach, combined with the ability of state-space models to capture complex dynamics and their interpretability, allows the FEU formalism for representing and analyzing neural data to serve as a tool to help bridge translation across model systems. Future work will aim to use FEDs to train real-time classifiers that can decode social representations from distributed population activity. Lastly, time series data occur widely in biological systems, and accurate unsupervised clustering can inform critical classification decisions. In conclusion, the FEU approach is generalizable and can provide utility in broad areas of biology, such as patient response data, genotyping, and diagnostic categorization.

## Methods

Details of electrophysiological data collection are found in Allsop et al. 2018 and Dal Monte et al. 2022.

### Simulating neural population activity

In order to test the accuracy of our clustering algorithm, we ran a simulation of 50 neurons categorized into five ground-truth clusters. Each cluster had a collection of 10 neurons that varied in their stimulus-response (i.e., excited/inhibited/no response) and sustainability of that response (i.e., sustained/phasic) in response to a stimulus. The five stimulus response groups were:

1. Excited/sustained - These neurons have an increased firing rate of +1 on a log scale in response to the stimulus.
2. Excited/phasic - These neurons have an increased firing rate of +1 on a log scale in response to the stimulus, but after a few time steps, the firing rate returns to what it was pre-stimulus.
3. Inhibited/sustained - These neurons have a decreased firing rate of −1 on a log scale in response to the stimulus.
4. Inhibited/phasic - These neurons have a decreased firing rate of −1 (on a log scale) in response to the stimulus, but after a few time steps, the firing rate returns to what it was pre-stimulus.
5. No response - These neurons have no change in firing rate relative to the stimulus.

To validate this pipeline more accurately test our pipeline, we injected noise into the simulation so that the change in firing rate was never exactly +1 or −1. Instead, it was the desired change + delta, where the delta is drawn from a Gaussian distribution with a standard deviation of 0.05. Thus, no two neurons had the same firing rate function.

### Comparing FEU pipeline to other clustering methods

We benchmarked our clustering algorithm against K-Means, T-SNE, PCA, and CEBRA to evaluate how the FEU analysis pipeline compares to existing methods. We simulated five datasets, each with a random number of neurons chosen between 30 and 50. Each simulation had five ground-truth clusters, but the number of data points in each cluster was different, varying in their stimulus response and sustainability of that response in response to a stimulus. The stimulus groups outlined in the previous section were excited/sustained, excited/phasic, inhibited/sustain, inhibited/phasic, and no response. Similar to the simulation in the preceding section, we injected noise into the simulation so that the change in firing was never exactly +1 or −1.

Since T-SNE, PCA, and CEBRA can be described as dimensionality reduction techniques and not clustering techniques, we combined these methods with K-Means to perform clustering analysis. This is done using K-Means clustering on the principal components or dimensions generated by the dimensionality reduction techniques. PCA was evaluated on 2, 5, and 10 principal components, while T-SNE and CEBRA were evaluated on 2 and 3 dimensions. In total, there were 9 combinations of techniques, firstly K-Means, followed by combinations of T-SNE, PCA, and CEBRA with K-Means, and finally, FEU analysis. For all tests, we assigned K-Means a cluster count of 5, which was the correct number of clusters in each simulation, reducing the task to only correctly assigning neurons to the right clusters, excluding the task of correctly predicting the number of clusters present in the data. However, being a non-parametric approach, FEU analysis will have a more difficult problem performing both tasks. Furthermore, we quantitatively evaluated the performance of these methods with statistical metrics, namely, normalized mutual information score (NMI), adjusted rand index (ARI), and accuracy. Accuracy in this context is the percentage of correctly predicted labels after the alignment of cluster labels between the true and predicted labels using the Hungarian algorithm.

### Analyzing electrophysiology data with the Functional Encoding Unit pipeline

Data from Allsop et al. 2018 and Dal Monte et al. 2022 was pre-processed and formatted into .csv files with identified time stamps of significant events (cue onset, face gaze onset, etc.) and the time stamps of all action potentials recorded from each neuron during each experiment. A stimulus (i.e., cue onset, face gaze onset, etc.) was selected to determine what FEU stimulus representation would be analyzed. Response windows were set to define the time window used to determine ensemble dynamics. For cue analysis during social learning, the baseline window was 500ms to 0ms, and the response window was 0ms to 1500ms. For face and object analysis during primate social gaze, a baseline window of 500ms to 0ms, and the response window was from 0ms to 500ms. These windows reflected the range of known single-neuron responses during the relevant stimulus and behavior (Allsop et al., Dal Monte et al.).

Python script was written to cluster neurons from a region or set of regions given their response to a defined stimulus during a defined response window. The algorithm defined by Lin et al. clusters neurons and provides the number of clusters as well as the number of neurons in each cluster and the values for the parameters (Jump, Phasicity) that define the cluster. The ‘Jump’ parameter refers to how the neuron cluster responds to the cue on a log scale. A positive Jump indicates that the firing rate increases, while a negative Jump indicates that the firing rate decreases. For example, if a cluster’s Jump is +1.2, all neurons in the cluster experience a firing rate multiplicative increase of e^1.2. The ‘Phasicity’ parameter is a normalized parameter that refers to the variability of the response during the response window interval after the desired stimulus. This is also on a log scale, and numbers are only interpretable relative to one another (for a given experiment, Phasicity is a random variable ‘var’ + a scalar constant =15).

Output from the code, available on GitHub, generated an Excel file with a Functional Encoding Dictionary detailing the number of Functional Encoding Unit (FEU) clusters, the number of neurons in each FEU, and parameter values for J and P. Code also generated graphs that plot each FEU ensemble as a circle in a 2D FED space in which the centroid of the circle represents its parameter values. The size of each circle represents the relative number of neurons in each FEU.

FEU rasters are generated as composite rasters made from individual neuron raster overlays evenly distributed along the y-axis. For display purposes, FEU rasters with large numbers of neurons use a random sampling of a subset of neurons to build composite rasters.

### Performing clustering analysis on electrophysiological data with benchmark methods

We applied K-Means and CEBRA + K-Means to the social learning data from Allsop et al. 2018, as discussed in the preceding section, to test their clustering analysis and ability to identify neural ensembles. We used a K-Means cluster count of 9, equivalent to the number of ensembles identified by FEU analysis on the same dataset. We show the resulting ensembles and display rasters of neurons assigned to each ensemble to evaluate the sanity of the predicted ensembles are shown.

### Mathematical framework for clustering algorithm

Consider an experiment with R successive trials, during which we record the activity of N neuronal spiking units. Let (0, T] be the continuous observation interval following the delivery of an exogenous stimulus at time τ = 0. We denote by {ΔN^(n)^}^T,R^ the discrete-time process obtained t,r t=1,r=1 by counting the number of events in T = ⌊T/Δ⌋ disjoint bins of width Δ = M ·δ, where M ∈ N. Given x^(n)^ and ψ^(n)^, the point process state-space model encodes the rate function {λ^(n)^}^T t=1^ of neuron n within an autoregressive process that underlies binomial observations. Applying the PPSSM independently to each neuron n from a population is akin to assuming that each neuron n is associated with a set of parameters θ, its own FEU. The goal of the framework introduced by Lin et al. is to group the N neurons into K FEUs, such that K is much smaller than N. To develop some intuition towards the time-series clustering model and algorithm from Lin et al. that lets us identify FEUs, suppose we knew the number K of FEUs. We will drop this assumption subsequently. Let z^(n)^ ∈ {1, · · ·, K} be the variable that denotes the index of the FEU to which neuron n belongs, and θ^(k)^ = [μ^(k)^, log ψ^(k)^]^⊤^ be the parameters that describe FEU k, k = 1, · · ·, K. We can obtain a clustered version of the PPSSM by assuming that neurons (n) that belong to the same FEU share the same set of parameters, i.e., θ such that z = k.

We can infer the number of FEUs can be inferred by using a so-called Chinese Restaurant Prior (CRP)^80,81^ on the FEU assignments and the number of FEUs^24^. Under a CRP, one (any) of the neurons first starts a FEU. Then, a second neuron either picks the same FEU as the first or starts a new FEU with some probability. This process is repeated such that every neuron picks an existing FEU with a probability proportional to the number of neurons already in that FEU or starts a new FEU with some probability. This process leads to anywhere between 1 and N FEUs, i.e., it needs not to assume that the number of FEUs is known as a priori. The goal is to infer, from the population activity, the number of FEUs (i.e., K), the parameters of each FEU µ, ψ), and the FEU to which each neuron n belongs (i.e. *Z*^(n)^). The full model is given by:

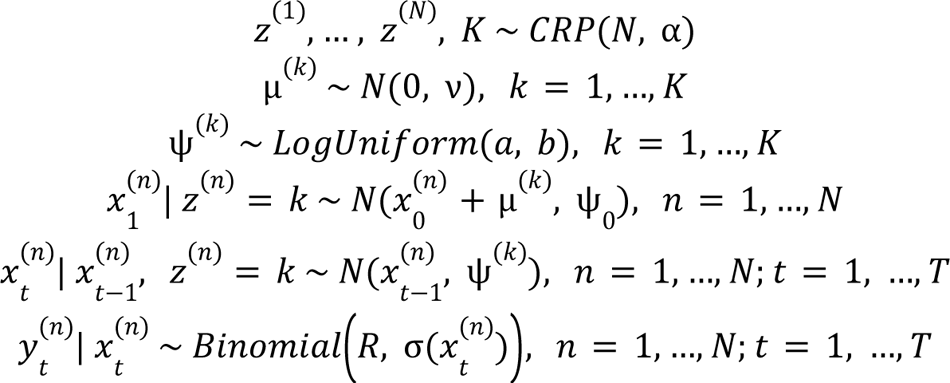

Here, *y^(n)^ = {y^(n)^_1_, …, y^(n)^_T_}* is a time series of observed firing sequences for neuron n. The PPSSM hypothesizes that each results from a latent, continuous process *x^(n)^ = {x^(n)^_1_, …, x^(n)^_T_}* that determines the unobserved firing probability over time, as calculated through the sigmoid function 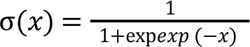. Neurons within a particular FEU k are assumed to share parameters that govern this latent process: The parameter μ^(k)^ describes the extent to which an exogenous stimulus modulates the response of the neurons within a cluster – a positive value of μ^(k)^ indicates excitation, a negative value indicates inhibition and a value close to zero indicates no response. We call this parameter the Jump (J) of the FEU. The state transition density f imposes a stochastic smoothness constraint on the rate function of neurons within a cluster, where ψ^(k)^ controls the degree of smoothness. A small value of ψ^(k)^ suggests that the neurons exhibit a sustained change in response to the stimulus, whereas a large value of ψ^(k)^ indicates that the change is unsustained, defined by. We can use this parameter as the Phasicity (P) of the FEU. The above-described process for identifying FEUs is automatic and unbiased in that it does not require any input from the experimentalist beyond the specification of the model and time window for analysis.

We also make some computational improvements to the inference algorithm of Lin et al. by adopting a stick-breaking representation of the CRP Chinese restaurant process; we enabled parallelizing the inference of the cluster assignments across different neurons, whereas Lin et al. had to infer these quantities sequentially per neuron. For datasets with a large number of neurons, the new algorithm is able to run much faster on multi-core hardware. Additionally, we provide an implementation of the algorithm that is compatible with graphics processing units (GPUs), which allows for further speedups. Our code is deposited at https://github.com/al5250/feu.

## Acknowledgments

We would like to thank Dr. Chris Pittenger for conceptual considerations regarding translation and research guidance during the development of this project. We thank Dr. Marina Picciotto for feedback on the FEU formalism and for making it more relevant to neuroscientists. We also thank Dr. Kafui Dzirasa for feedback on FEU formalism and machine learning approaches in neuroscience. AA, DB conceptualized and developed the FEU framework; AL, DB derived computational algorithms; AA collected rodent data; ODM, SF collected primate data; KT supervised rodent experiment design and data collection; SC supervised primate experiment design and data collection; AA and AL organized and pre-processed rodent data; NF and PP organized and pre-processed primate data; AL, AA, CA performed FEU analysis; CA performed benchmarking experiments. AA wrote the manuscript; AA, AL, CA, DB, SC, and KT edited it. The rodent and computational work was supported by T32MH014276 and T32MH019961 grants, the National Institute of Mental Health, Yale Department of Psychiatry, Burroughs Wellcome Fund Collaborative Research Travel Grant (AA), an NDSEG Fellowship (AL), R37-MH102441 and HHMI Investigator Program (KMT). The primate work was supported by NIMH R01MH110750 and NIMH R01MH120081(SC).

## Data Accessibility

All data is accessible, and the code is available on GitHub.

