## Supplementary Materials for "An unbiased method to partition diverse neuronal responses into functional ensembles reveals interpretable population dynamics during innate social behavior"

### Supplementary information

### Table 1 Functional Encoding Dictionary for data in Figure 5

| ACC | | | | | | | | |
|---|---|---|---|---|---|---|---|---|
| social learning cue | | | | | | | | |
| habituation | | | | | conditioning | | | |
| FEU | Jump | Phasicity | count (n=195) | | | jump | Phasicity | count (n=195) |
| 1 | 1.013753 | 9.874331 | 45 | | 1 | 0.708874 | 9.267389 | 89 |
| 2 | 0.256558 | 0.544152 | 14 | | 2 | 1.254519 | 9.952948 | 28 |
| 3 | 0.380936 | 9.887316 | 6 | | 3 | 0.118707 | 5.706698 | 20 |
| 4 | -0.178783 | 2.294918 | 16 | | 4 | -0.153754 | 9.562564 | 22 |
| 5 | 1.742927 | 9.671714 | 14 | | 5 | 0.234562 | 8.000342 | 21 |
| 6 | 0.511013 | 8.512954 | 39 | | 6 | -0.722206 | 9.014569 | 12 |
| 7 | 0.036585 | 8.940191 | 27 | | 7 | 2.396977 | 9.810637 | 2 |
| 8 | -0.337037 | 0.514732 | 6 | | 8 | -2.566214 | 5.372104 | 1 |
| 9 | -0.722206 | 9.014569 | 21 | | | | | |
| 10 | -2.051888 | 8.544649 | 2 | | | | | |
| 11 | 0.485651 | 0.806981 | 5 | | | | | |

| ACC | | | | | | | | |
|---|---|---|---|---|---|---|---|---|
| control analysis | | | | | | | | |
| habituation | | | | | conditioning | | | |
| FEU | Jump | Phasicity | count (n=195) | | | Jump | Phasicity | count (n=195) |
| 1 | -0.100197 | 3.992866 | 69 | | 1 | -0.154046 | 1.878003 | 46 |
| 2 | 0.415732 | 0.449714 | 17 | | 2 | 0.012521 | 4.616255 | 95 |
| 3 | 0.14902 | 0.180749 | 56 | | 3 | 0.27781 | 5.170693 | 36 |
| 4 | 0.094573 | 9.836461 | 14 | | 4 | 0.808602 | 8.957752 | 10 |
| 5 | -0.255283 | 7.643664 | 13 | | 5 | -0.412457 | 2.676885 | 8 |
| 6 | -0.722206 | 9.014569 | 14 | | | | | |
| 7 | -0.479157 | 1.868132 | 9 | | | | | |
| 8 | 1.852387 | 5.133327 | 1 | | | | | |
| 9 | 1.851931 | 4.899814 | 1 | | | | | |
| 10 | -2.525573 | 0.290565 | 1 | | | | | |

**Table 2 Functional Encoding Dictionary for data in Figure 6**

| ACC | | | | | | |
|---|---|---|---|---|---|---|
| habituation | | | | | | |
| FEU | Jump | Phasicity | count (n=195) | photoidentified % | excited network % | inhibited network % |
| 1 | 1.013753 | 9.874331 | 45 | 35.7 | 21.4 | 6.6 |
| 2 | 0.256558 | 0.544152 | 14 | 7.1 | 14.3 | 6.6 |
| 3 | 0.380936 | 9.887316 | 6 | 7.1 | 14.3 | |
| 4 | -0.178783 | 2.294918 | 16 | | 3.6 | 20 |
| 5 | 1.742927 | 9.671714 | 14 | 7.1 | 17.9 | |
| 6 | 0.511013 | 8.512954 | 39 | 14.3 | 17.9 | 26.7 |
| 7 | 0.036585 | 8.940191 | 27 | 14.3 | 7.1 | |
| 8 | -0.337037 | 0.514732 | 6 | | | 13.3 |
| 9 | -0.722206 | 9.014569 | 21 | 14.3 | | 26.7 |
| 10 | -2.051888 | 8.544649 | 2 | | | |
| 11 | 0.485651 | 0.806981 | 5 | | 3.5 | |
| ACC | | | | | | |
| conditioning | | | | | | |
| FEU | Jump | Phasicity | count (n=195) | photoidentified % | excited network % | inhibited network % |
| 1 | 0.708874 | 9.267389 | 89 | 64.3 | 71.4 | 40 |
| 2 | 1.254519 | 9.952948 | 28 | 14.3 | 10.7 | |
| 3 | 0.118707 | 5.706698 | 20 | | 7.1 | |
| 4 | -0.153754 | 9.562564 | 22 | 14.3 | 3.6 | 26.6 |
| 5 | 0.234562 | 8.000342 | 21 | | 3.6 | 20 |
| 6 | -0.722206 | 9.014569 | 12 | 7.1 | | 13.3 |
| 7 | 2.396977 | 9.810637 | 2 | | 3.6 | |
| 8 | -2.566214 | 5.372104 | 1 | | | |

**Table 3 Functional Encoding Dictionary for data in Figure 7**

| ACC face | | | | | ACC object | | | |
|---|---|---|---|---|---|---|---|---|
| FEU | Jump | Phasicity | count (n=236) | | FEU | Jump | Phasicity | count (n=184) |
| 1 | -0.015117 | 0.171617 | 172 | | 1 | 0.019435 | 3.489979 | 156 |
| 2 | 0.213257 | 5.90104 | 18 | | 2 | -1.836143 | 8.44577 | 2 |
| 3 | -0.162247 | 7.371203 | 18 | | 3 | -0.24348 | 4.779605 | 15 |
| 4 | -0.233201 | 6.88877 | 23 | | 4 | 0.232237 | 3.772021 | 7 |
| 5 | 0.130502 | 2.9514 | 5 | | 5 | -0.747206 | 7.171842 | 1 |
| | | | | | 6 | 0.351108 | 8.117762 | 3 |
| BLA face | | | | | BLA object | | | |
| FEU | Jump | Phasicity | count (n=537) | | FEU | Jump | Phasicity | count (n=393) |
| 1 | 0.074587 | 1.003158 | 91 | | 1 | -0.050213 | 5.800307 | 63 |
| 2 | -0.01666 | 0.904745 | 225 | | 2 | -0.004533 | 0.676951 | 88 |
| 3 | 0.023357 | 7.41292 | 63 | | 3 | -0.129041 | 3.837597 | 52 |
| 4 | -0.169746 | 9.043318 | 35 | | 4 | 0.063474 | 0.886419 | 72 |
| 5 | -0.20652 | 2.821232 | 25 | | 5 | 0.108349 | 7.725325 | 56 |
| 6 | 0.166382 | 5.056808 | 38 | | 6 | -1.534201 | 4.325586 | 4 |
| 7 | 0.024505 | 4.674267 | 51 | | 7 | -0.027084 | 6.219159 | 29 |
| 8 | 0.419689 | 9.682577 | 6 | | 8 | -0.13327 | 9.843607 | 29 |
| 9 | 0.176288 | 7.079611 | 1 | | | | | |
| 10 | 1.23387 | 1.581351 | 2 | | | | | |
| dmPFC face | | | | | dmPFC object | | | |
| FEU | Jump | Phasicity | count (n=187) | | FEU | Jump | Phasicity | count (n=139) |
| 1 | -0.017626 | 2.024209 | 108 | | 1 | 0.242413 | 9.723268 | 6 |
| 2 | -0.092583 | 1.096741 | 19 | | 2 | 0.021292 | 6.405596 | 103 |
| 3 | -0.096001 | 8.671568 | 3 | | 3 | -0.258845 | 7.160438 | 29 |
| 4 | 0.109218 | 1.214583 | 22 | | 4 | -0.455267 | 0.312573 | 1 |
| 5 | 0.053151 | 8.77568 | 26 | | | | | |
| 6 | 0.004324 | 8.82068 | 9 | | | | | |
| OFC face | | | | | OFC object | | | |
| FEU | Jump | Phasicity | count (n=241) | | FEU | Jump | Phasicity | count (n=195) |
| 1 | -0.238121 | 9.936711 | 10 | | 1 | -0.044021 | 8.474085 | 57 |
| 2 | 0.047788 | 7.135459 | 76 | | 2 | 0.008295 | 3.102671 | 51 |
| 3 | -0.01392 | 3.259947 | 68 | | 3 | 0.118643 | 4.917292 | 15 |
| 4 | -0.097605 | 5.022757 | 38 | | 4 | 0.086658 | 5.138108 | 26 |
| 5 | -0.066333 | 7.810266 | 49 | | 5 | -0.081089 | 2.374053 | 45 |
| | | | | | 6 | -1.451036 | 0.653735 | 1 |

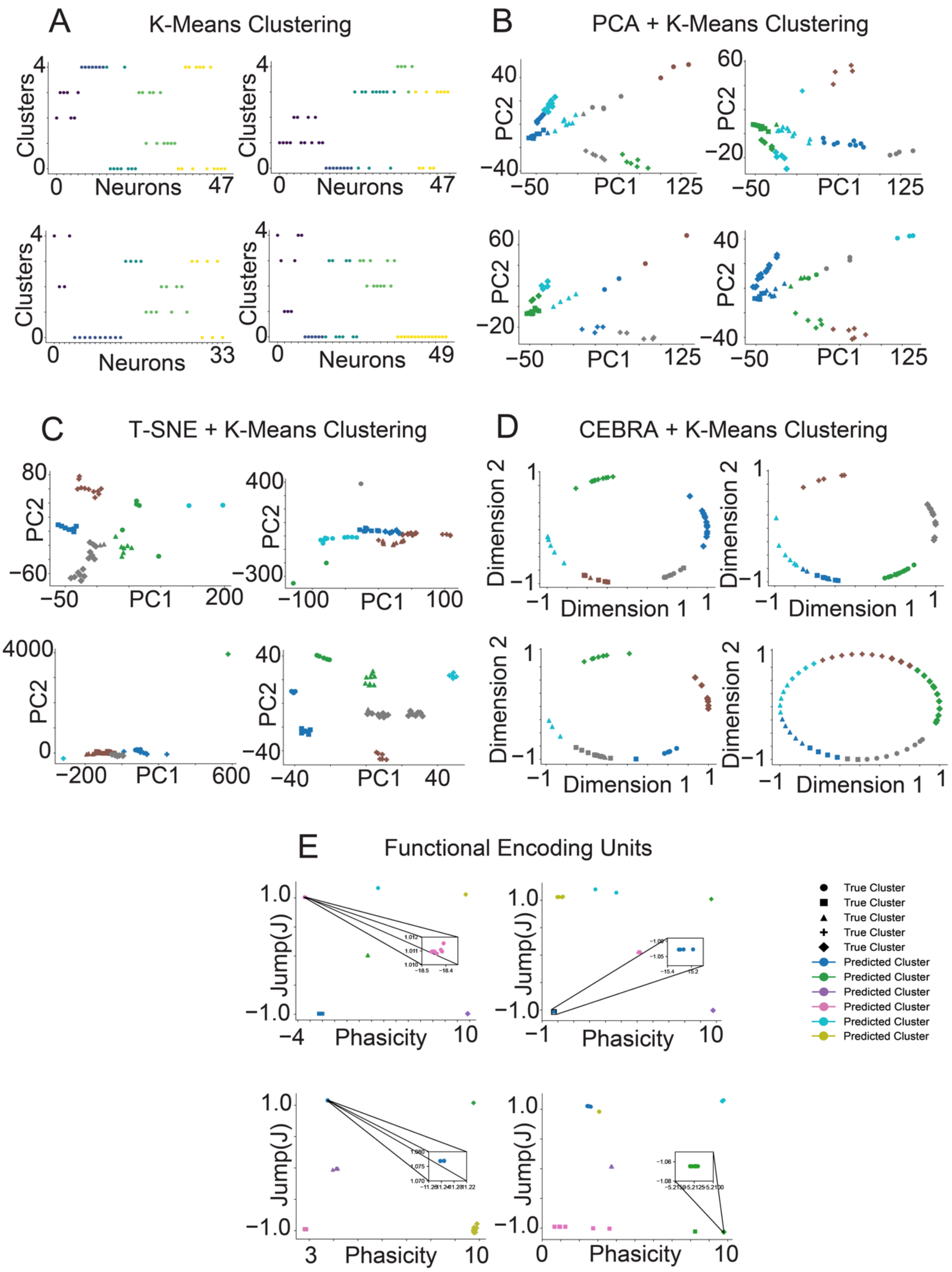

A
K-Means Clustering
B
PCA + K-Means Clustering
C
T-SNE + K-Means Clustering
D
CEBRA + K-Means Clustering
E
Functional Encoding Units
Clusters
Neurons
PC1
PC2
Dimension 1
Dimension 2
Jump(J)
Phasicity
True Cluster
Predicted Cluster

**Supplemental Figure 1**

**FEU pipeline, on average clusters simulated data more accurately than other methods**

We apply the 5 techniques to cluster 5 simulated datasets into ensembles. Across the 5 experiments (experiment 1 is shown in Figure 3), the FEU pipeline obtains the best average performance. This is evident in the visualizations of these clusters.

**A.** We applied K-Means to cluster simulated neuron spike data. Data points correspond to individual neurons; the y-axis 'Clusters' denotes each neuron's cluster assignments determined by K-Means, with colors indicating the neurons' true clusters. Perfect clustering is achieved when neurons within the same predicted cluster share identical colors, reflecting accurate alignment with their true cluster.

**B.** We performed dimensionality reduction on the simulated neuron spike data with PCA and clustered the resulting principal components with K-Means. Data points correspond to individual neurons; data point shape denotes the ground truth cluster assignment, while colors denote the predicted clusters. Perfect clustering is achieved when data points in clusters have the same color and shape.

**C.** We conducted dimensionality reduction on the simulated neuronal spike data using T-SNE, followed by clustering the derived principal components with K-Means. Each data point represents an individual neuron, with the shape of the data point indicating the neuron's actual cluster assignment and colors representing the predicted clusters. Perfect clustering is achieved when data points in clusters have the same color and shape.

**D.** We performed dimensionality reduction on the simulated neuronal spike data utilizing CEBRA and subsequently clustered the resultant components with K-Means. Each data point signifies an individual neuron; data point shape denotes the ground truth cluster assignment, while colors denote the predicted clusters. Perfect clustering is achieved when data points in clusters have the same color and shape.

**E.** FED representation of the simulated neuron spike data. We use FEU analysis to cluster the simulated neuron spike data. Perfect clustering is achieved when data points in each cluster have the same color and shape. Callout to show overlapping data points in a cluster.

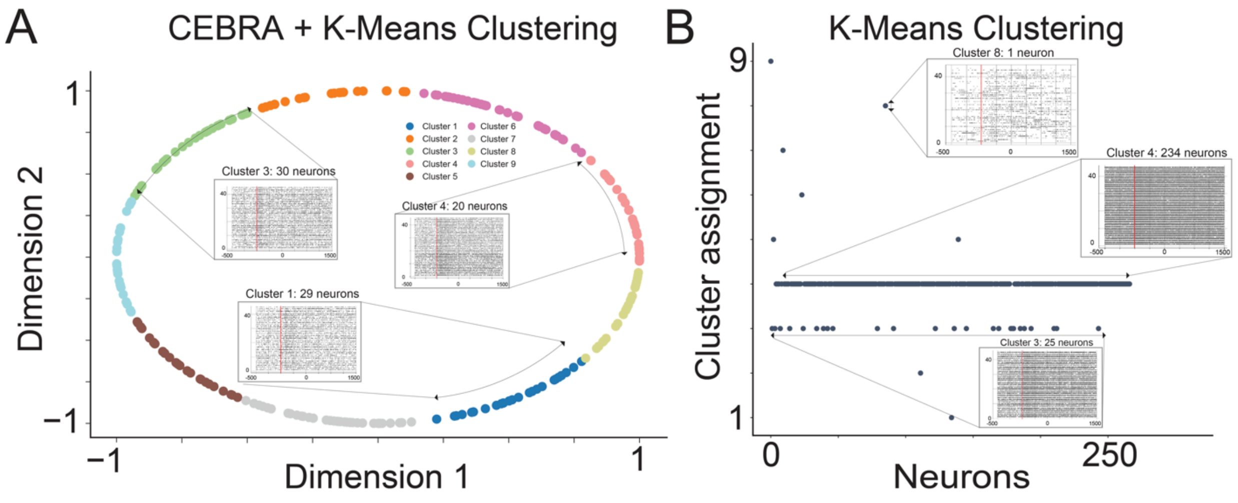

**Supplemental Figure 2**

**Alternative methods do not efficiently capture ensembles in real trial data**

**A.** We use CEBRA to perform dimensionality reduction and K-Means to cluster the neural spike data from the rodent social learning tasks.

**B.** We use K-Means to cluster neural spike data from the rodent social learning experiment.

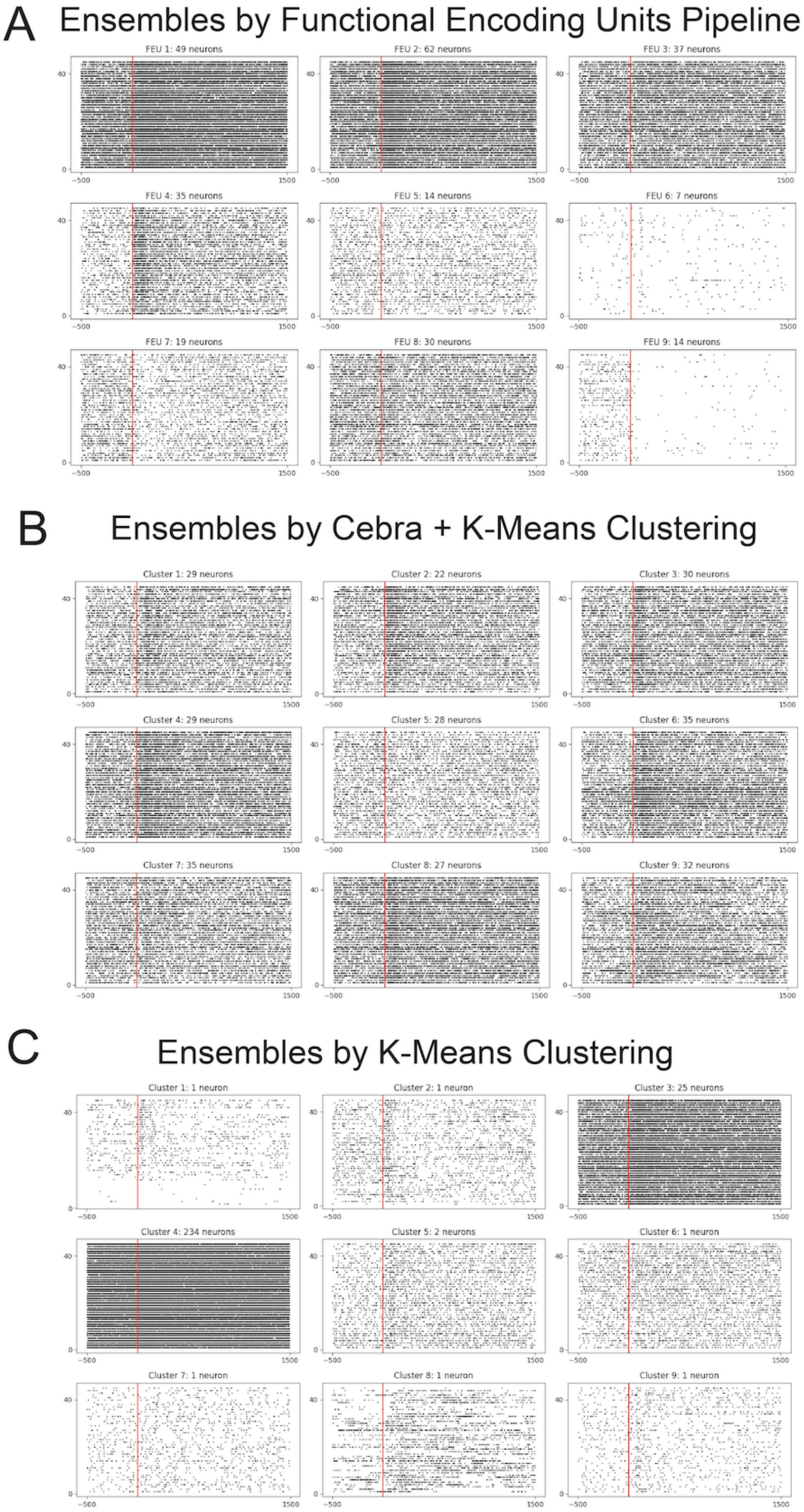
A
Ensembles by Functional Encoding Units Pipeline
FEU 1: 49 neurons
FEU 2: 62 neurons
FEU 3: 37 neurons
FEU 4: 35 neurons
FEU 5: 14 neurons
FEU 6: 7 neurons
FEU 7: 19 neurons
FEU 8: 30 neurons
FEU 9: 14 neurons
B
Ensembles by Cebra + K-Means Clustering
Cluster 1: 29 neurons
Cluster 2: 22 neurons
Cluster 3: 30 neurons
Cluster 4: 29 neurons
Cluster 5: 28 neurons
Cluster 6: 35 neurons
Cluster 7: 35 neurons
Cluster 8: 27 neurons
Cluster 9: 32 neurons
C
Ensembles by K-Means Clustering
Cluster 1: 1 neuron
Cluster 2: 1 neuron
Cluster 3: 25 neurons
Cluster 4: 234 neurons
Cluster 5: 2 neurons
Cluster 6: 1 neuron
Cluster 7: 1 neuron
Cluster 8: 1 neuron
Cluster 9: 1 neuron

**Supplemental Figure 3**

**Alternative methods do not efficiently capture ensembles in real trial data**

A. Ensembles found by the FEU pipeline.

B. Ensembles predicted by the CEBRA + K-Means approach of clustering demonstrated limited interpretability.

C. Ensemble predicted by the K-Means clustering technique shows a significant lack of substantive coherence.
